# Roles for RERE in lymphatic endothelial cell proliferation and survival, and human cystic lymphatic malformations

**DOI:** 10.1101/2024.03.24.586490

**Authors:** Daniella M. Rogerson, Ajit Muley, Jessica Giordano, Zoe Vogel, Ronald Wapner, Carrie J. Shawber

## Abstract

Human congenital lymphatic anomalies (LAs) arise due to defects in lymphatic development. During a genetic study of fetuses with LAs, we identified a heterozygous pathogenic truncating variant in *RERE* in a fetus with a cystic lymphatic malformation (CLM). RERE is a transcriptional regulator which interacts with several key lymphangiogenic factors, including Notch and Coup-TFII. RERE also modulates retinoic acid signaling, which is essential for lymphatic vascular development. Thus, we hypothesized that RERE functions in lymphatic endothelial cells (LECs) and its loss contributes to LEC dysfunction and CLM pathogenesis. RERE was found to be expressed in the lymphatic endothelium during human development. *RERE* knockdown in human LECs reduced proliferation and induced apoptosis, increased expression of key lymphangiogenic genes, *PROX1, COUP-TFII* and *VEGFR3*, and altered expression of Notch target genes. RERE expression was elevated in LECs isolated from CLMs with pathogenic *PIK3CA* variants. These findings support a novel role for RERE in LECs, where RERE regulates LEC proliferation, LEC survival, lymphangiogenic gene expression and Notch signaling, which in turn suggests its loss contributes to CLM pathogenesis.

## INTRODUCTION

The lymphatic vasculature serves many essential functions, including returning interstitial fluid to the circulatory system, transporting immune cells and signaling molecules, and absorbing lipids from the intestine (1). In humans, congenital lymphatic anomalies (LAs) arise due to defects in development of the lymphatic vasculature, and include cystic lymphatic malformations (CLMs), generalized lymphatic anomalies (GLAs), central collecting lymphatic anomaly (CCLA), and primary lymphedema. LAs often arise and are identified in utero, however, diagnosing LAs remains challenging, especially in the prenatal period (2, 3). An understanding of the genetic etiologies of LA is required to allow for diagnosis and clinical interventions.

The arginine-glutamic acid dipeptide repeats gene, *RERE* (*ATROPHIN 2*), encodes a nuclear receptor modulator capable of interacting with numerous epigenetic regulators in mice and humans via its N-terminal ELM2 and SANT domains (4-6). In humans, *de novo* germline heterozygous variants in *RERE* are associated with Neurodevelopmental Disorder with or without Anomalies of the Brain, Eye, or Heart (NEDBEH) (7). Individuals with NEDBEH present with visual and hearing impairments, intellectual disability, central nervous system abnormalities, congenital heart disease, genitourinary anomalies, and non-specific dysmorphic facial features. In mice, RERE is essential for development, and *Rere*-null mice parallel human phenotypes, with cardiac defects, CNS abnormalities, asymmetric somitogenesis, and optic and CNS abnormalities that lead to embryonic death due to cardiac failure shortly after E9.5 (8).

Although *Rere*-null mice die prior to lymphatic development, RERE has been shown to interact with key regulators of lymphangiogenesis including retinoic acid (RA) and Coup-TFII. RERE is a positive modulator of RA signaling in the mouse embryo, where RERE is recruited as part of a complex with Coup-TFII to promote RA target gene expression (8, 9) (10). Coup-TFII is itself essential for murine lymphangiogenesis, where it promotes LEC specification via induction of and interaction with *Prox1*, and its loss in lymphatic endothelium disrupts lymphatic development (11, 12) (13) (14) (15). RA signaling is an important regulator of early lymphangiogenesis. In murine embryoid bodies, treatment with RA promoted LEC differentiation, while treatment with a retinoic acid receptor (RAR) inhibitor suppressed specification (16). *In vitro*, ectopic 9-cis RA promoted LEC proliferation and migration, and *in vivo* induced lymphatic sprouting and vessel formation (17). In murine development, excess RA signaling led to a dysfunctional lymphatic vasculature that was hyperplastic, hemorrhagic, and associated with edema (18). Conversely, reduced RA signaling led to the development of hypoplastic lymphatic vasculature, and edema (18) (19). Thus, both an excess or deficiency of RA signaling has been associated with impaired lymphangiogenesis and lymphatic dysfunction.

RERE has also been suggested to positively modulate Notch signaling, which has been shown to suppress LEC specification and is necessary for valve formation during murine development (14, 20-28). RERE was identified as part of a Notch transcriptional activator complex (28). In the chick neural tube, *Rere* loss-of-function (LOF) led to reduced expression of the Notch effectors, *Hes* genes, while RERE overexpression upregulated Notch target gene expression (28). Coup-TFII and Notch also regulate each other via complex feedback loops. COUP-TFII suppresses Notch target gene expression directly, and NOTCH signaling in turn negatively regulates *Coup-tfII* expression (15, 21). As RERE interacts with RA signaling, *COUP-TFII*, and *NOTCH* signaling, all of which in concert are modulators of lymphatic endothelial development, we hypothesized that RERE functions in LECs and its loss contributes to lymphatic defects and dysfunction.

Here, we describe a *de novo* heterozygous frameshift variant in *RERE* predicted to be LOF in a 9-week fetus presenting with a CLM (29). We found that RERE was expressed in embryonic and postnatal lymphatic endothelium during human development. *In vitro*, RERE was necessary for LEC proliferation and survival, and its loss led to increased expression of several key lymphangiogenic genes including *PROX1, COUP-TFII* and *VEGFR3*, and altered expression of Notch target genes. Lastly, *RERE* expression was elevated in LECs isolated from CLMs, especially CLMs carrying *PIK3CA* variants. These data support a novel role for RERE in LEC proliferation, survival, suppression of lymphangiogenic gene expression, which in turn may contribute to CLM pathogenesis.

## MATERIAL AND METHODS

### Clinical samples and whole exome sequencing (WES)

Neonatal foreskins, fetal tissues, and LA specimens from biopsy and resection were collected from December 11, 2010 to May 16, 2017 as approved by the Columbia Institutional Review Board (AAAO8009, AAAI0082, AAAA7338, AAAA9976) and studies using these specimens or cells derived from these specimens were performed between May 31, 2022. Authors had access to information that could identify individual participants during and after data collection for a subset of samples collected. LECs were isolated from LAs as previously described (30). LECs and parental and fetal samples from case Fetal0045F were subjected to WES at the Columbia University Institute for Genomic Medicine using the NimblegenSeqCap EZ V2.0/3.0, SeqCap EZ HGSC VCRome, or xGenExome Research Panel v1.0 kits with the Illumina NovaSeq platform (29). Reads were aligned using human reference GRCh37 with Illumina DRAGEN Bio-IT Platform, and duplicates were marked with Picard (Broad Institute). Variants were called using Genome Analysis Toolkit Best Practices v3.6 and annotated with ClinEff and Analysis Tool for Annotated Variants (in-house Analysis Tool for Annotated Variants) with American College of Medical Genetics (ACMG) criteria. All variants were confirmed by sanger sequencing (29).

### Tissue immunostaining

Tissues were fixed in 4% paraformaldehyde (PFA) overnight at 4°C, equilibrated in 30% sucrose in 1×PBS at 4°C, and embedded in OCT. Fixed frozen sections (5 μm) were post-fixed in cold acetone, washed 2 times with 1×PBS, blocked with 2% donkey serum, 3% BSA in 1×PBS for 1 hour at room temperature and then incubated with an anti-RERE antibody and anti-human PODOPLANIN overnight 4°C (**Supplementary Table 1**). The following day primary antibodies were detected with Alexa Fluor–conjugated secondary antibodies as previously described (2). RERE and PODOPLANIN antibody specificity was determined by comparing tissue stained with secondary antibodies only (**Supplemental Figure 1**). Images were captured with Olympus IX83 fluorescence microscope with Olympus CellSense software. RERE positive lymphatics were scored as PODOPLANIN positive vessels with either continuous or discontinuous RERE staining.

### Lymphatic endothelial cell culture and lentiviral transduction

Human dermal lymphatic endothelial cells (HdLECs; PromoCell) were cultured on fibronectin-coated plates in EBM-2 Basal Medium with EGM-2 MV supplements (Lonza) and 10 ug/mL VEGF-C (RnD Systems). Three lots of HdLECs were used. HdLECs were lentivirally infected with pLKO plasmid encoding either scrambled sequence (SCR) or *RERE* specific shRNA, TRCN0000329758 (Sigma-Aldrich) (31). After 72-96 hours post transfection, or 72-96 hours of puromycin selection (2ug/mL), *RERE* knockdown was confirmed by qRT-PCR and cells from five independent transductions were utilized for experiments with data presented for at least three independent transductions (**primers in Supplement Table 2**).

### Cell immunofluorescence

HdLECs were seeded on fibronectin-coated 4-well Millicell EZ Slides (MilliporeSigma). After either 24- or 48-hours, cells were washed with cold 1×PBS, fixed in 4% PFA for 15 minutes on ice, washed 3 times in cold 1×PBS and incubated in blocking solution (5% donkey serum, 0.1% Triton X-100 in 1×PBS) for 1 hour at room temperature (RT) followed by an overnight 4°C incubation with an antibody against KI67 or RERE (**Supplementary Table 1**). Negative controls were incubated in blocking solution without primary antibody. The following day, slides were washed 3 times with cold 1×PBS and incubated with Alexa Fluor–conjugated secondary antibodies (**Supplementary Table 1**) for one hour at RT, washed 3 times with 1×PBS and coverslipped with mounting media with DAPI (Vector). Images were captured with Zeiss Axioskop2 Plus/Zeiss AxioCam MRc camera with Zen software (RERE) or Olympus IX83 fluorescence microscope with Olympus CellSense software (KI67). For KI67, at least eight images per experiment and condition were analyzed with ImageJ and the number of Ki67+ nuclei were determined as a percentage of total DAPI+ nuclei per slide. Data presented for at least three independent experiments.

### Cell TUNEL Assay

Lentivirally infected HdLECs were seeded on fibronectin-coated 4-well Millicell EZ Slides (MilliporeSigma). 24 hours later cells were washed 3 times in cold 1×PBS, fixed in 4% PFA for one hour on ice and permeabilized in 0.1% Triton X-100 in 0.1% sodium citrate for two minutes on ice. Apoptosis was determined using the *In Situ* Cell Death Detection Kit (Sigma-Aldrich) per manufacture’s protocol. For negative controls, slides were incubated in label solution only without enzyme. Slides were washed 3 times in cold 1×PBS and coverslipped with mounting media with DAPI. Images were captured with Olympus IX83 fluorescence microscope with Olympus CellSense software. At least 6 images per experiment and condition were analyzed with ImageJ and the number of TUNEL+ nuclei were determined as a percentage of total DAPI+ nuclei per slide. Data presented for at least three independent experiments.

### Semiquantitative and quantitative RT-PCR

RNA was extracted using the NucleoSpin RNA Mini kit (Machenery Nagel) or the RNeasy Mini Kit (Qiagen). 1-2 ug of total RNA was DNaseI (Ambion) digested followed by first strand syntheses using random hexamers and Superscript II reverse transcriptase (Invitrogen). A list of primers for which amplicons were confirmed for specificity by Sanger sequencing are presented in **Supplementary Table 2**. For semiquantitative RT-PCR, cDNA was amplified with platinum Taq polymerase (Invitrogen) for *RERE* (30 cycles) and *β-ACTIN* (20 cycles) using a thermocycler (Eppendorf) and then separated on an agarose gel (Alkali Scientific). For qRT-PCR, serially diluted matching human PCR amplicons cloned into the pDrive vector (Qiagen) were used to generate a standard curve. PCR reactions were prepared in triplicate with Sybr Green Master Mix (Applied Biosystems) and amplified using a CFX96 PCR Cycler (Bio-rad). Data was normalized by *β-ACTIN* levels. Data presented for at least three independent experiments.

### Statistical Analysis

Data were analyzed with Prism9. Means were compared by Unpaired t-test or Welch’s t-test in the setting of a large difference in standard deviation, or by Ordinary One-way ANOVA or Welch’s and Brown-Forsythe ANOVA with multiple comparisons. A p-value ≤ 0.05 was considered significant.

## RESULTS

### A LOF *RERE* variant was identified in a 9-week fetus with a CLM

Review of a previously published cohort of fetal anomalies in which parental/fetal trios underwent whole exome sequencing (WES) identified a heterozygous *RERE* truncation variant in a fetus diagnosed with a 3.5 mm septated CLM at the posterior nuchal region in the first trimester (9w6d; Fig 1A) (7, 29). The variant was predicted to cause haploinsufficiency via early truncation of the *RERE* protein four amino acids downstream of the frameshift site. There were no microarray findings in this case; a concurrent novel VOUS in *EPHB4* was identified (29). Though the CLM appeared by imaging to resolve in the second trimester, a persistent nuchal fold consistent with a CLM was diagnosed postnatally. The neonate was noted to have dysmorphic features, recurrent urinary tract infections, and spasticity and hypertonia on neurological exam, consistent with NEBDEH. Notably, cardiac abnormalities were absent in this case. Whether the CLM persisted or continued to grow or whether developmental delay consistent with NEBDEH presented pediatrically is unclear, as the patient was lost to follow-up at 8 months of age (7, 32).

**Figure 1.**
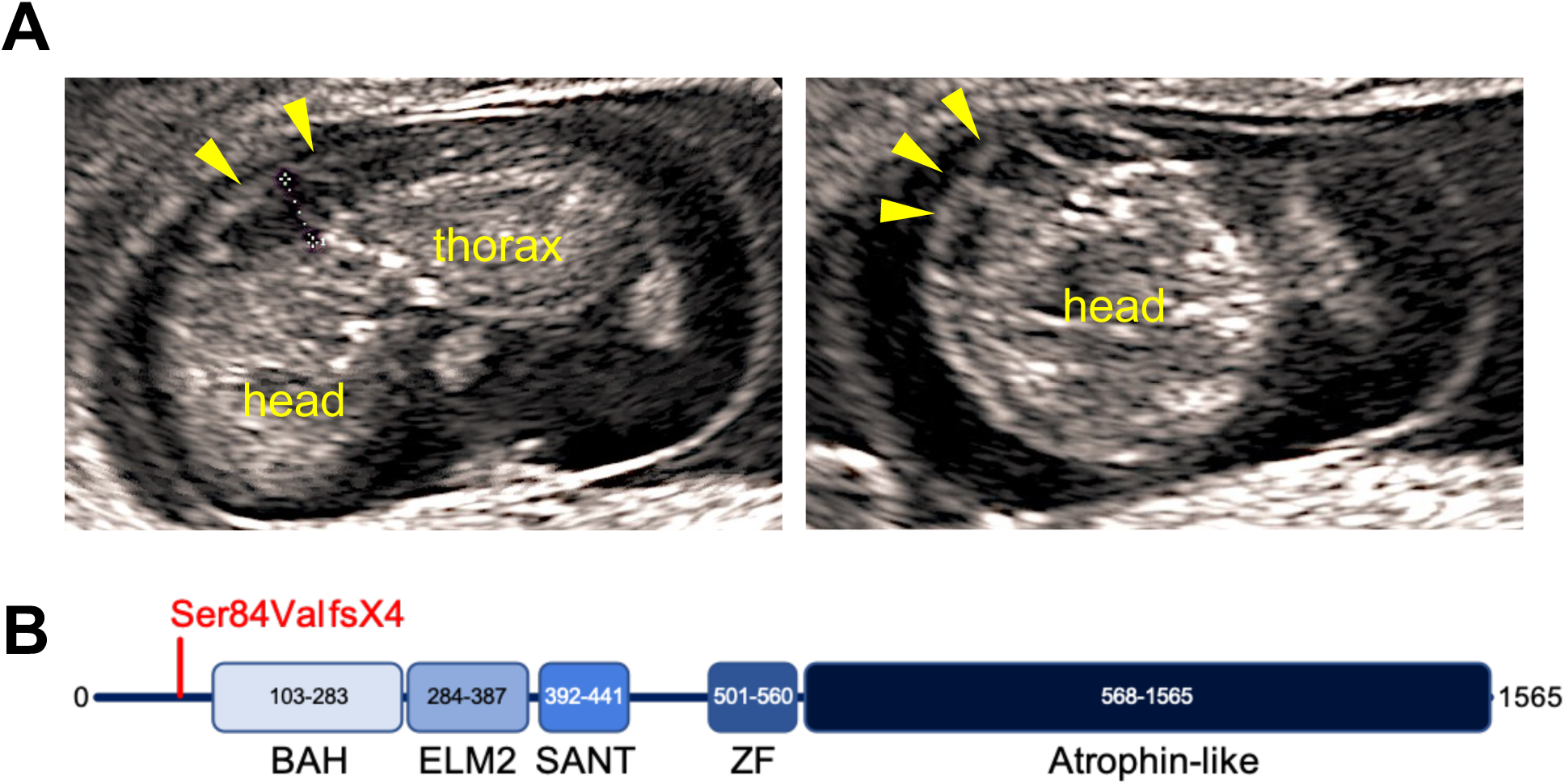
Pathogenic *RERE* LOF variant in a fetal CLM. **A)** Ultrasound (sagittal left, axial right) of a dorsal septated 3.5 mm CLM at the neck (yellow arrows and dashed caliper) of a 9-week and 6-day of gestation (9w6d) fetus carrying a *RERE* LOF variant. **B**) Diagram of RERE protein structure. A pathogenic *de novo* heterozygous germline frameshift variant (red text) in *RERE* was identified by WES. BAH – Bromo adjacent homology domain, ELM2 – Egl-27 and MTA1 homology 2 domain, SANT – SANT/Myb domain, ZF – GATA type zinc finger domain, Atrophin-like – atrophin-like domain.

In this case, a germline *de novo* heterozygous c.248dupA single nucleotide insertion variant in *RERE* was identified. This frameshift variant, p.Ser84fsX4, is absent from large population databases, falls within the first exon of the canonical transcript, and is predicted to lead to early termination of the protein four amino acids downstream of the variant, precluding translation of any RERE functional domains (**Fig 1B**). As *RERE* is sensitive to haploinsufficiency, and LOF is a known mechanism of disease, the RERE p.Ser84fsX4 variant was classified as pathogenic for NEBDEH per ACMG criteria and likely causative of the patient’s phenotype (32). However, pathogenic variants in *RERE* have not previously been associated with CLMs, nor has a role for RERE in the lymphatic endothelium been described.

### Human fetal and neonatal lymphatic endothelial cells express RERE

To determine the expression of RERE in human LECs during development, fetal nuchal tissues from second trimester human fetal specimens (15-21 weeks gestational age) and neonatal dermis (day of life 2) were immunostained for RERE and the LEC protein, PODOPLANIN. RERE was expressed in the endothelium of a subset lymphatic and blood vessels from 16 to 21 weeks of gestation, ranging between 25% and 47% of lymphatic vessels assessed **(Fig 2A)**. No trend in RERE expression over fetal development was noted, though the small sample size of lymphatic vessels at each gestational age limited analyses. In the neonatal dermal lymphatic vasculature, RERE expression was continuously or partially expressed in the lymphatic endothelium of 14.81% (4 of 27) of vessels, as well as a subset of blood vessels **(Fig 2B)**. Analysis of neonatal HdLEC in culture demonstrated that RERE was ubiquitously expressed in a punctate manner in the nucleus and cytoplasm **(Fig 2C)**. To determine relative expression of *RERE* in the blood and lymphatic endothlelial cells, *RERE* expression in HdLECs and human umbilical venous endothelial cells (HUVECs) was determined by semi-quantitative RT-PCR **(Fig 2D)**. Both HdLECs and HUVECs expressed *RERE*, but its expression was higher in HUVECs. Together, these data demonstrate that RERE is expressed in a subset of human LECs during fetal development supporting a role for RERE in lymphatic endothelium.

**Figure 2.**
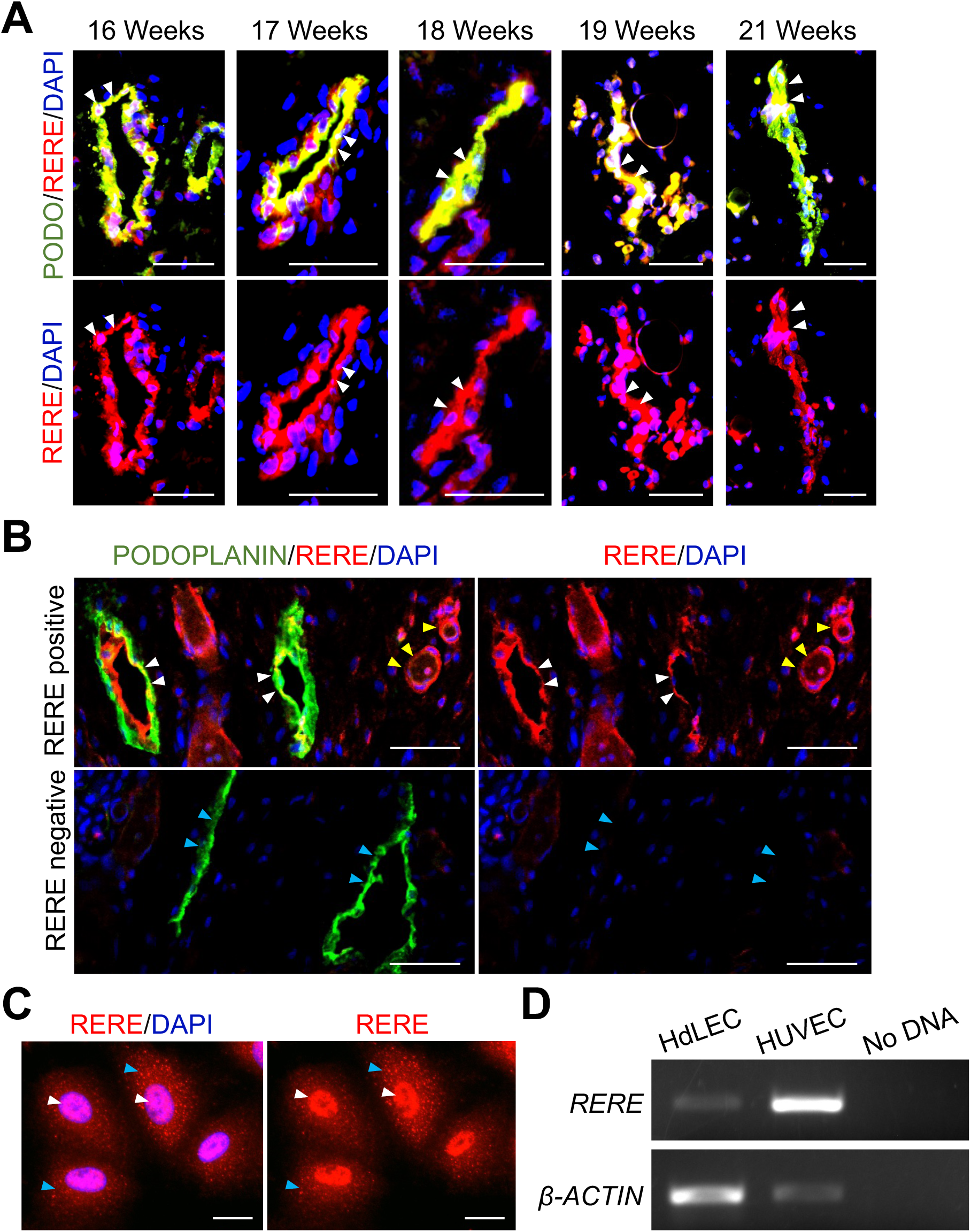
RERE was expressed in human fetal and neonatal lymphatic endothelium. **A)** 15-to 21-week fetal tissue through the dorsal neck region (n=9) and **B)** 2-day old neonatal skin was stained for RERE and the lymphatic protein, PODOPLANIN; white arrowhead – RERE positive lymphatic vessels, blue arrowhead – RERE negative lymphatic vessel, yellow arrowhead – blood vessel. Scale bars: 50 μM. **C)** HdLECs were stained for RERE; white arrowhead – nuclear staining, blue arrowhead – cytoplasmic punctate staining. Scale bars: 20 μM. **D)** *RERE* (30 cycles) and *β-ACTIN* (20 cycles) RT-PCR of HdLECs and human umbilical venous endothelial cells (HUVECs) compared to a no DNA control.

### RERE was necessary for LEC proliferation and survival

To determine the role of RERE in human LECs, we knocked-down *RERE* in HdLECs using a lentivirus encoding an *RERE* shRNA (sh*RERE*) and compared findings HdLECs infected with a lentivirus containing a scrambled sequence (SCR). qRT-PCR confirmed *RERE* was knocked down approximately 50% relative to controls which mimicked the heterozygous LOF variant identified in the fetal CLM **(Fig 3 A)**. To determine the effects of *RERE* knockdown on proliferation, HdLECs were stained for the proliferative marker, KI67. sh*RERE* HdLEC proliferation was reduced 20% relative to SCR HdLECs **(Fig 3 B-C)**. sh*RERE* HdLECs also developed stress granules consistent with increased cell death. To quantify apoptosis, TUNEL staining was performed. Apoptosis was significantly increased 3-fold in sh*RERE* HdLECs relative to SCR controls **(Fig 3 D-E)**. Together, these data suggest the RERE functions to promote LEC viability and proliferation.

**Figure 3.**
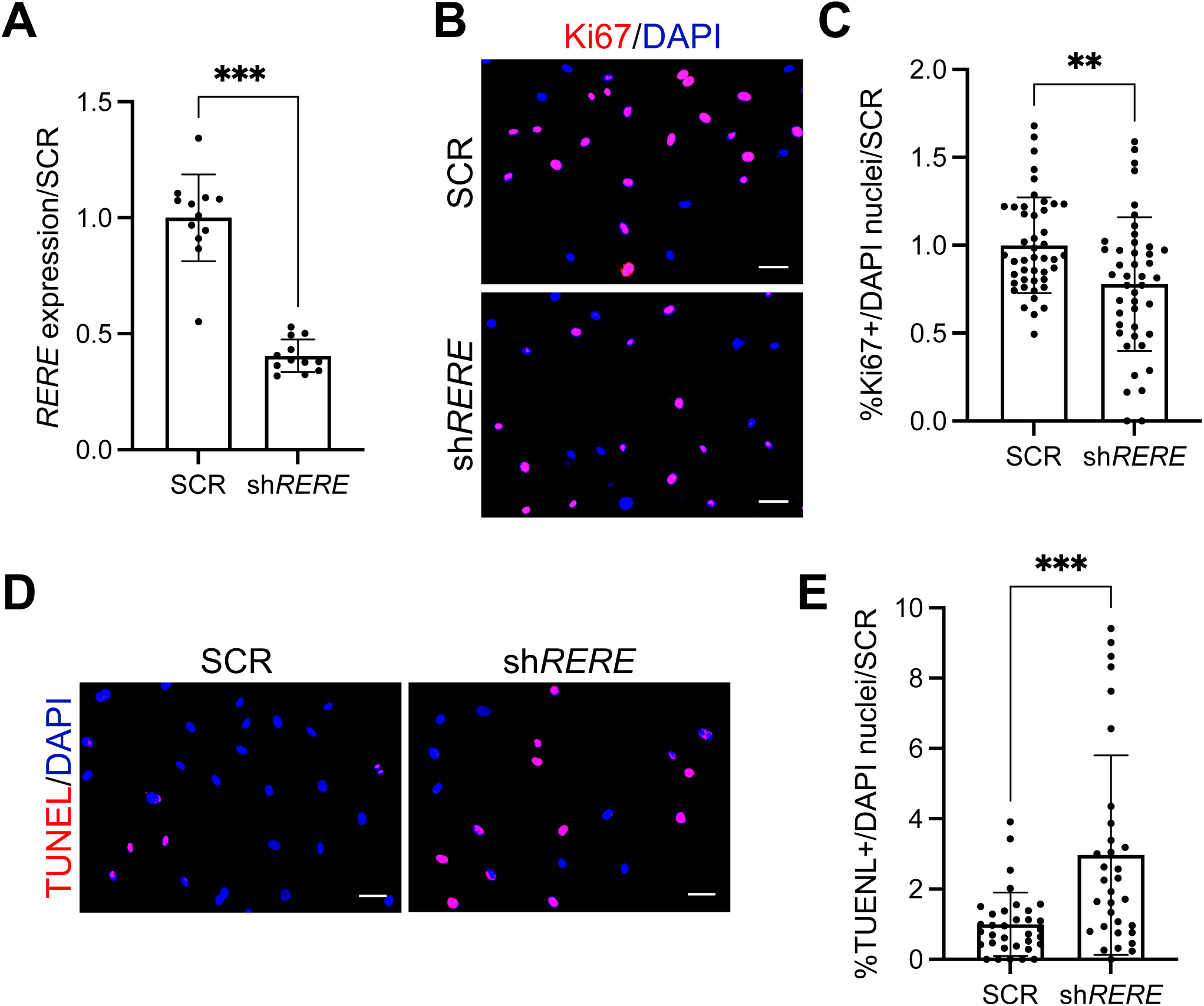
RERE suppressed HdLEC proliferation and apoptosis. **A)** *RERE* qRT-PCR of scrambled control (SCR) and *RERE* knockdown (sh*RERE*) HdLECs. *RERE* expression normalized to *β-ACTIN* expression and presented relative to SCR control ± STD.***p<0.0001. **B)** sh*RERE* and SCR HdLECs stained for KI67. Scale bars 50μM. **C)** Quantification of KI67 expression presented as percent KI67+ nuclei normalized to total nuclei per field of view relative to SCR control ± STD. *p=0.0022. **D)** TUNEL performed on sh*RERE* and SCR HdLECs. Scale bars 50 μM. **E)** Quantification of TUNEL presented as TUNEL+ nuclei normalized to total nuclei per field of view relative to SCR control ± STD. **p=0.0006.

### RERE regulated transcription of expression of lymphangiogenic genes and Notch effectors

As RERE is a nuclear receptor modulator which has been shown to regulate gene expression, we determined expression of a panel of lymphangiogenic genes in sh*RERE* HdLEC and SCR HdLECs (8) (4-6). Expression of the pro-lymphangiogenic genes *PROX1, COUP-TFII*, and *VEGFR3* were significantly increased in sh*RERE* HdLECs relative to SCR HdLECs **(Fig 4 A-C)**. As RERE has been shown to modulate *NOTCH* signaling and downstream gene expression, we also determined the expression of the *NOTCH* effectors, *HES1, HEY1* and *HEY2*. Expression of *HEY1* was significantly reduced, while *HEY2* expression was increased in sh*RERE* HdLECs relative to controls **(Fig 4 D-E)**. Expression of *HES1* was unchanged. Together, these data demonstrate a role for RERE in suppressing pro-lymphangiogenic genes and modulating NOTCH effectors in LECs.

**Figure 4.**
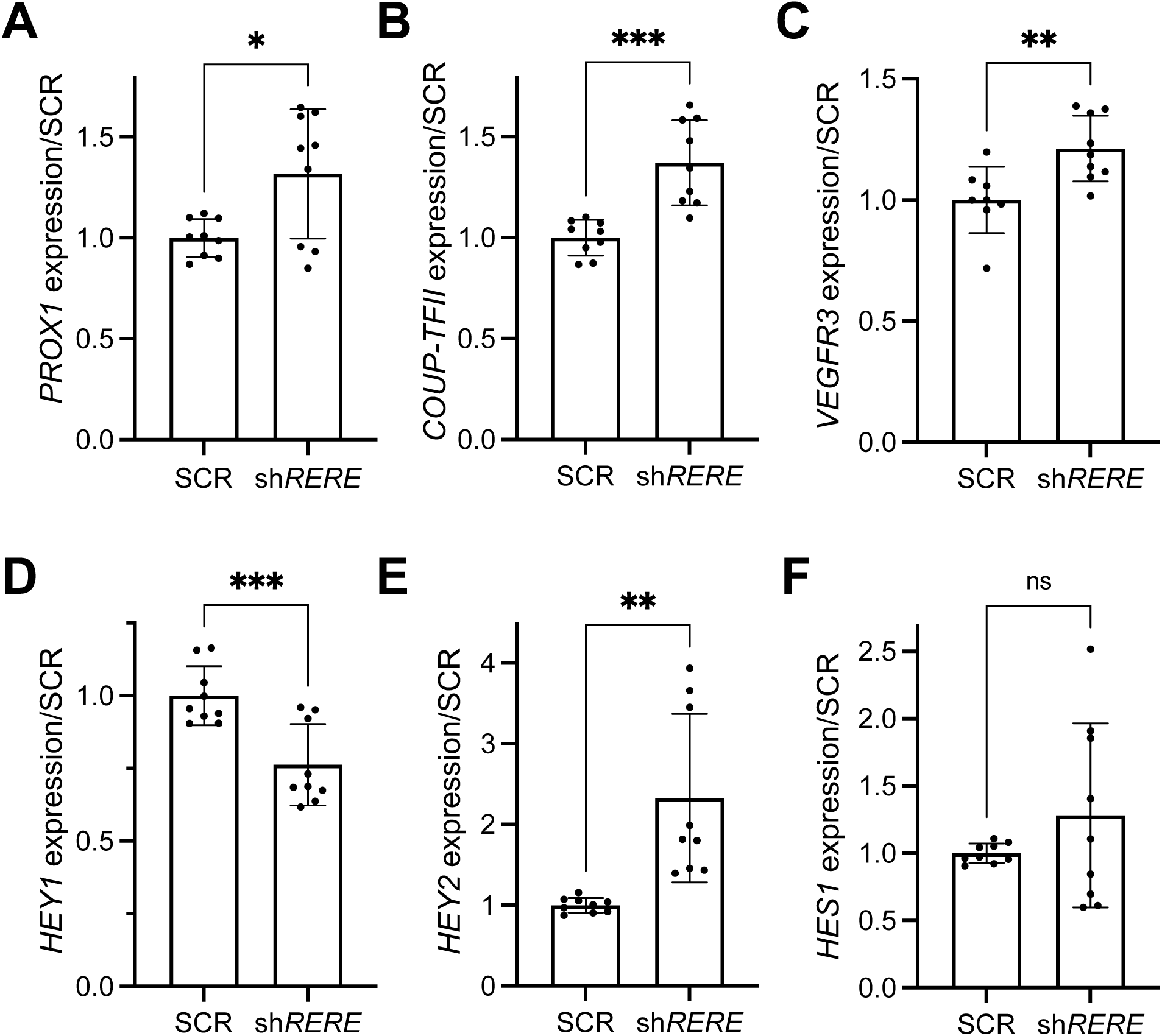
Lymphangiogenic and Notch effector gene expression were altered in HdLECs with *RERE* knockdown. qRT-PCR of *RERE* shRNA (sh*RERE*) and scrambled control (SCR) HdLECs for **A)** *PROX1*, **B)** *COUPTFII*, **C)** *VEGFR3*, **D)** *HEY1*, **E)** *HEY2*, and **F)** *HES1* normalized by *β-ACTIN* relative to SCR ± STD. ns – not significant, *p<0.02, **p<0.006, *** p<0.0009.

### *RERE* transcripts were elevated in LECs isolated from CLMs with *PIK3CA* variants

As *RERE* haploinsufficiency was seen in a human case of CLM, we determined *RERE* expression via qRT-PCR in a cohort of LECs isolated from human LAs (n=21) for which *PIK3CA* variant status was determined. RERE expression was elevated in LECs from CLMs (n=15) relative to HdLEC controls (n=3). In contrast, *RERE* expression was similar in LECs from GLA (n=1) and CCLA (n=5) to HdLECs **(Fig 5 A)**. Of the 15 CLM included in this cohort, 7 LECs carried pathogenic variants in *PIK3CA. RERE* expression was elevated in LECs with *PIK3CA* variants compared to HdLEC controls, as well as compared LECs from CLMs without *PIK3CA* variants **(Fig 5 B)**. Among LECs with *PIK3CA* variants, *RERE* expression was highest in LECs with variants within the kinase domain compared to variants in the helical domain **(Fig 5C)**. These data demonstrate that *RERE* is upregulated in LECs isolated from CLMs and especially among LECs from CLMs with activating *PIK3CA* variants.

**Figure 5.**
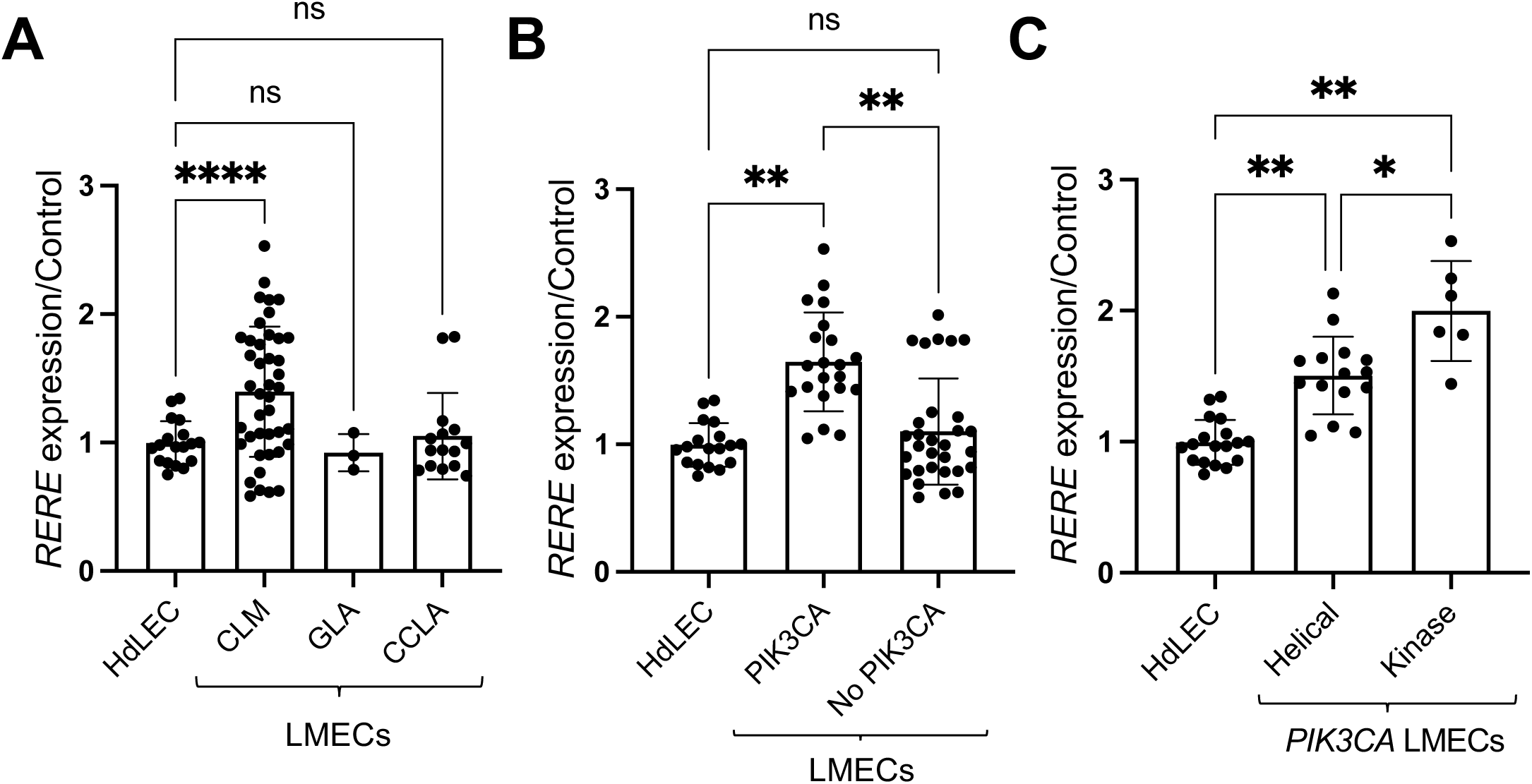
*RERE* expression was increased in LECs from CLMs carrying *PIK3CA* variants. qRT-PCR of a cohort of LECs (n=21) isolated from CLM (n=15), GLA (n=1) and CCLA (n=5) relative to HdLEC controls (n=3). **A)** *RERE* expression compared between LECs from LAs and control HdLECs **B)** *RERE* expression in CLMs (n=17) compared between those with *PIK3CA* variants (n=7) relative to those without *PIK3CA* variants (n=10). **C)** *RERE* expression in LECs from CLMs with *PIK3CA* variant analyzed by affected domain. Helical – helical domain (n=6), Kinase – kinase domain (n=2). Data presented in triplicate normalized to *β-ACTIN* expression relative to HdLEC controls ± STD. ns – not significant, *p=0.0011, **p<0.0001.

## DISCUSSION

We identified a pathogenic *RERE* LOF variant in a fetus with CLM suggesting RERE functions lymphatic development and anomalies. Expression studies showed that RERE was expressed in the lymphatic endothelium of a subset of vessels during human development and in growing neonatal HdLECs. *RERE* haploinsufficiency in HdLECs was associated with reduced proliferation and increased cell death. RERE also functioned to suppressed lymphangiogenic gene expression, and modulated expression of the NOTCH effectors, *HEY1* and *HEY2*. Finally, *RERE* transcripts were elevated in LECs isolated from CLMs, especially those that carry *PIK3CA* variants. Together, these data demonstrate that RERE promoted HdLEC proliferation and survival, suppressed lymphangiogenic gene expression, and dynamically modulated Notch target gene expression, supporting a role for *RERE* in lymphatic development and possibly its dysregulation in the development of CLM.

RERE is a known modulator of RA signaling, where RERE is recruited as part of a complex with COUP-TFII and RAR at RA response elements of target genes (9, 10). The pathogenic variant described in this case is predicted to lead to early truncation of the *RERE* protein, precluding expression of the N-terminal ELM2 and SANT domains, which are important for binding with transcriptional regulators (4-6). The truncated RERE protein in the fetal CLM may thus be unable to interact within the transcriptional complex to modulate RA signaling. *RERE* haploinsufficiency was associated with reduced proliferation, and RA signaling has been shown to promote lymphatic differentiation and proliferation (17). The converse findings of *RERE* haploinsufficiency in the case of fetal CLM, but increased *RERE* expression in LECs isolated from CLMs, may be explained exquisite sensitivity of the lymphatic endothelium to RA signaling. This has been established in early murine lymphangiogenesis, where both too much and too little RA signaling were associated with aberrant lymphatic development, and in one model, nuchal edema (16) (19) (18). Thus, both an excess or deficiency in *RERE* may lead to anomalies in lymphatic development due to dysregulation of RA signaling.

*RERE* LOF led to increased expression of key lymphangiogenic genes, *COUP-TFII, VEGFR3* and *PROX1*, suggesting a suppressive role for RERE in LEC gene expression. Coup-TFII is necessary for early LEC specification via induction of Prox1, the master regulator of LEC specification (11, 12, 14) (15). VEGFR3 signaling promotes LEC specification and maintenance, as well as proliferation, and migration in a context dependent manner (13, 33). As Coup-TFIis a known co-activator of Prox1 which in turn induces *VEGFR3* expression in LECs, increased expression of *PROX1* and *VEGFR3* in *RERE* haploinsufficient HdLECs could be secondary to increased Coup-TFII expression (19) (34). The effect of the upregulation of these key lymphangiogenic genes after *RERE* LOF remains unknown. *RERE* suppression of the major lymphangiogenic genes *COUPFTII, PROX1* and *VEGFR3* implicates *RERE* in the most fundamental stages of lymphatic development.

RERE has been described to positively modulate Notch signaling and effector expression in neuronal cells, possibly by stabilization of the NICD and transactivation of the transcription factor RBPjK/CBF1 (28). Consistent with RERE having both repressive and positive regulatory transcriptional roles, RERE knockdown dynamically modulated NOTCH effector gene expression, where its loss led to downregulation of *HEY1* and upregulation of *HEY2* (8) (4-6). Notch signaling plays an established role in lymphangiogenesis, where it serves early in lymphatic development to suppress LEC specification, while later in development it is essential for lymphatic valve formation (14, 20-27). Differential expression of Notch effectors and lymphatic genes have also been described for Notch1 and Notch4 signaling (20). As Notch plays complex and highly context dependent roles in lymphangiogenesis and LEC proliferation and survival, it is possible that RERE modulation of Notch effectors contributed to the altered lymphangiogenic gene expression, LEC proliferation, and LEC viability we found in *RERE* deficient LECs.

This is the first reported case of a pathologic *RERE* variant implicated in a fetal CLM. The patient later developed mild neurological symptoms and genitourinary anomaly consistent with *RERE* associated NEBDEH. One case of NEBDEH due to an Atrophin-like domain missense variant in *RERE* has been described to have redundant nuchal skin on pediatric exam, whether this case represented CLM or was an incidental finding is unknown (7). The unique lymphatic presentation of this case of NEBDEH, with only mild neurological findings and absence of cardiac abnormalities, could be due to the truncating LOF variant observed in *RERE*, as individuals with point mutations in the commonly affected Atrophin-like domain of RERE have more severe NEBDEH phenotypes than their counterparts with LOF variants (7). There is significant overlap between neuronal, cardiac and lymphatic developmental genes, and RA is a well-established modulator of neuronal differentiation in addition to regulating the earliest stages of lymphangiogenesis (1, 19, 33, 35). Lymphatic endothelial cell dysfunction has also been described in patients with neonatal congenital heart defects and chylothorax (36). The fact that RERE functions have been implicated in lymphatic development, as well as neuronal development, may explain the combination of lymphatic and neurological phenotypes in this case of NEBDEH.

Expression studies of HdLECs also suggest a role for *RERE* in CLMs, as we found that *RERE* expression was elevated in CLM LECs, especially those that carried *PIK3CA* variants. The PI3K/AKT pathway is a major signaling network downstream of VEGFR3 activation in LECs responsible for survival, proliferation and cell cycle progression (34). Somatic activating variants in *PIK3CA* are pathogenic in human CLMs and GLAs, and are associated with LEC hyperproliferation and elevated expression of lymphangiogenic genes (37) (38) (39) (40) (41) (42). As we found that *RERE* LOF was associated with reduced proliferation and increased cell death, it is possible that *RERE* expression was increased in LMECs with *PIK3CA* variants as they are hyperproliferative (43, 44).

## CONCLUSIONS

We established that the nuclear receptor modulator, RERE, was expressed in human developing lymphatic endothelium and was upregulated among LECs from CLM. Data supports a role for RERE in promoting LEC proliferation and cell survival, suppressing lymphangiogenic gene expression, and modulating Notch signaling. These findings support the consideration of *RERE* gene evaluation in the setting of fetal CLM. How RERE modulates proliferation, apoptosis and gene expression in LECs is unknown. Further work examining RERE in lymphangiogenesis, LEC proliferation, and survival is needed, as is work characterizing a role for RERE a as a modulator of RA and *NOTCH* signaling.

## ACKNOWLEDGEMENTS

We thank Michael Schonning for providing images demonstrating specificity of the antibody against human PODOPLANIN and Dr. June K. Wu for IRB oversight for the collection and secondary use of LA specimens.

