## Supporting Data for "Roles for RERE in lymphatic endothelial cell proliferation and survival, and human cystic lymphatic malformations"

Supplementary Table 1: Antibody Information

| Antibody | Catalog Number | Supplier | Host | Clone | Antigen | Antibody Registry ID | Dilution | Validated |
|---|---|---|---|---|---|---|---|---|
| KI67 | ab15580 | Abcam | Rabbit | Polyclonal | Synthetic peptide. Ab15581 | AB_443209 | 1:200, Cell Immunofluorescence | By supplier |
| RERE | ab103462 | Abcam | Rabbit | Polyclonal | Human RERE amino acids 389-418 | AB_10710905 | 1:50, Cell & Tissue Immunofluorescence | Supplemental Figure 1A,B. |
| PODOPLANIN | AF3670 | R&D Systems | Sheep | Polyclonal | Human PODOPLANIN amino acids 21-123 | AB_2162070 | 1:200, Tissue Immunofluorescence | Supplemental Figure 1C. |
| Donkey anti-Sheep IgG Secondary Antibody, Alexa Fluor™ 488 | A11015 | Invitrogen | Donkey | Polyclonal | IgG (H+L) ovine | AB_2534082 | 1:1000, Tissue Immunofluorescence | By supplier |
| Donkey anti-Rabbit IgG Secondary Antibody, Alexa Fluor™ 594 | A21207 | Invitrogen | Donkey | Polyclonal | IgG (H+L) rabbit | AB_141637 | 1:1000 Cell & Tissue Immunofluorescence | By supplier |

Supplementary Table 2: RT-PCR Primers

| Gene | Forward Primer | Reverse Primer |
|---|---|---|
| *RERE* | 5' CCTGTAGGGAGTAAGAGGGACCATC 3' | 5' GGGTACTGTTCAGCCTCCTTGTC 3' |
| *B-ACTIN* | 5' CGAGGCCCAGAGCAAGAGAG 3' | 5' CTCGTAGATGGGCACAGTGTG 3' |
| *PROX1* | 5' ACGTAAAGTTCAACAGATGCATTAC 3' | 5' ACGTAAAGTTCAACAGATGCATTAC 3' |
| *COUP-TFII* | 5' GCCATAGTCCTGTTCACCTC 3′ | 5' CTGAGACTTTTCCTGCAAGC 3′ |
| *VEGFR3* | 5' GAGACCTGGCTGCTCGGAAC 3' | 5' GAGACCTGGCTGCTCGGAAC 3' |
| *HEY1* | 5' ACGAGAATGGAAACTTGAGTTC 3' | 5' AAC TCC GAT AGT CCA TAGCAAG 3' |
| *HEY2* | 5' ATGAGCATAGGATTCCGAGAGTG 3' | 5' GGCAGGAGGCACTTCTGAAG 3' |
| *HES1* | 5' CCCAACGCAGTGTCACCTTC 3' | 5' TACAAAGGCGCAATCCAATATG 3' |

Rogerson, et al. Supplemental Figure 1

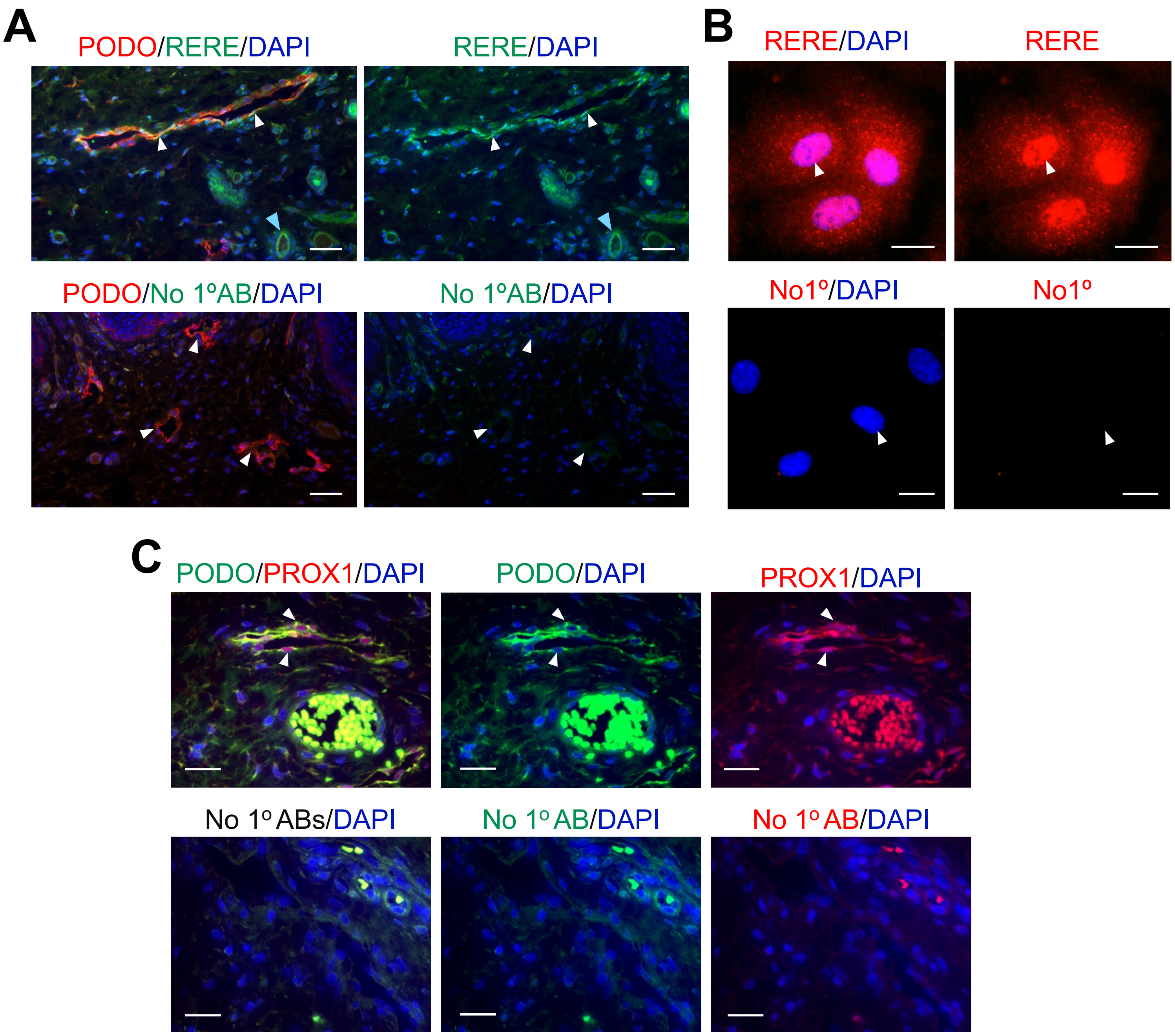


**Supplemental Figure 1. Validation of RERE and PODOPLANIN antibodies.**
**A)** Human neonatal skin stained for PODOPLANIN and RERE (top) versus PODOPLANIN and secondary antibody only (No 1°AB) for RERE (bottom). White arrowhead – lymphatic vessels. Blue arrowhead – blood vessels. Scale bars: 50 μM. **B)** HdLECs stained for RERE (top) or secondary antibody only (bottom). White arrowhead – nuclei. Scale bars: 20 μM. **C)** Human neonatal skin stained for human PODOPLANIN (PODO) and the lymphatic endothelial cell marker, PROX1 (top) versus stained with secondary antibodies only (No 1° AB; bottom). White arrowhead – lymphatic vessels. Scale bars: 50 μM.
